# Readout-dependent time-of-flight weighting of absorption and flow in interferometric diffuse optics

**DOI:** 10.64898/2026.09.02.748230

**Authors:** Marcin Marzejon, Neda Mogharari, Klaudia Nowacka-Pieszak, Dawid Borycki

## Abstract

Joint optical measurement of hemoglobin and blood-flow dynamics could provide a compact route to richer functional monitoring of tissue physiology. However, hybrid diffuse-optical systems commonly optimize photon time-of-flight (TOF) for individual readouts, even when absorption and flow are recovered from the same acquisition. Here, we asked whether a single TOF window can optimally preserve both signals. Dual-wavelength TOF-iSCOS recovered absorption, hemoglobin and relative blood-flow index (BFI) from the same interferometrically sensed optical field, while Early, Middle and Late photon windows provided progressively greater weighting of longer and deeper photon paths. During forearm cuff occlusion in 11 adults, absorption and flow showed markedly different TOF dependence: Late-gate BFI responses were retained, whereas absorption decreased to a median Late/Early ratio of 0.18. The resulting readout-by-TOF interaction was positive in all 11 participants (median 2.61 octaves, 95% bootstrap CI 2.44–3.22; exact sign-test P = 9.8 × 10⁻⁴). Readout-specific gate selection increased held-out response retention by 24.8% (95% CI 8.8–42.5%) relative to a single common gate. We then tested the approach during prefrontal working-memory activation in seven participants using a higher-load 2-back task relative to a low-load 0-back control. Late-gate BFI increased in all seven participants (+5.07%, 95% CI 1.46–8.68%; P = 0.014) and remained positive after adjustment using an Early-TOF signal, whereas hemoglobin responses were less consistent. Layered Monte Carlo modeling showed that the same detected photon histories acquire different absorption and dynamic-scattering sensitivity across TOF, linking the experimental effects to anatomy- and geometry-conditioned tissue weighting. Within the present geometry and photon budget, the central result is therefore not that one readout is intrinsically superior, but that absorption and flow recovered from the same optical field retain physiological information differently across photon time-of-flight.

## 1. Introduction

Diffuse optical neurophysiology rests on two complementary measurements. Intensity-based near-infrared spectroscopy (NIRS) uses wavelength-dependent attenuation and the modified Beer–Lambert law (MBLL) to estimate changes in oxy-, deoxy- and total hemoglobin, whereas diffuse correlation spectroscopy (DCS) and related field-decorrelation methods infer microvascular blood-flow changes from the temporal decorrelation of multiply scattered light [1–4]. These observables report different parts of the hemodynamic response: flow regulates oxygen delivery, whereas hemoglobin concentrations reflect blood volume, venous drainage and oxygen extraction. Their coupling is neither linear nor instantaneous. Flow changes can be larger and earlier than volume- and oxygenation-based responses [5–9]. Hybrid NIRS–DCS systems therefore combine the two readouts to monitor flow, oxygenation and oxidative metabolism in muscle and brain [10–14]. Measuring both is what separates oxygen delivery from its downstream vascular and metabolic consequences.

The two contrasts also do not weight tissue identically. Conventional intensity-based functional NIRS at adult source–detector separations are strongly affected by extracerebral absorption and task-locked scalp hemodynamics [15–18]. Flow-sensitive methods are not immune to superficial contamination, but their brain-to-scalp sensitivity is predicted and observed to exceed that of continuous-wave NIRS at matched geometry, because cerebral perfusion differs more strongly from scalp perfusion than cerebral hemoglobin concentration differs from its extracerebral counterpart [13,19]. A measured difference between hemoglobin and flow responses may therefore reflect both vascular physiology and the photon populations that carry each contrast. Existing NIRS–DCS combinations establish the value of joint measurement, but separate intensity and correlation channels cannot by themselves reveal whether the same photon time-of-flight (TOF) window is optimal for both.

Time-resolved diffuse optics adds photon TOF as a control over optical pathlength and depth sensitivity [20–24]. Later-arriving photons travel longer mean paths and carry greater deep-tissue weighting, yet their flux falls sharply in the temporal tail, and their estimates become more sensitive to the instrument response function (IRF), gate width, source–detector separation, motion and model error [24,25]. TOF gates are overlapping sensitivity distributions rather than isolated anatomical layers, so an optimal gate must balance depth weighting against photon statistics. TD-DCS gate optimization has accordingly been formulated as a trade-off among deep sensitivity, photon statistics, superficial rejection and IRF characteristics [26,27]: later is not universally better. Crucially, intensity changes and field decorrelation are different functionals of the same detected field, so increasing TOF need not improve absorption-derived hemoglobin and flow estimates by the same amount.

Interferometric near-infrared spectroscopy provides a uniquely direct way to test this proposition. Swept-source interferometry recovers the complex optical field as a function of photon TOF [28]. The gate-integrated field magnitude supplies wavelength-resolved intensity for MBLL, while temporal statistics of the same field within the same gates supply the first-order field autocorrelation *g*_1_ and finite-exposure speckle visibility *k*^2^ used to recover BFI [23,29,30]. MBLL’s direct gate integration, intensity log-ratio and pathlength scaling underpin fNIRS, but whether this simple reduction preserves functional contrast in the low-support temporal tail is unresolved. A single acquisition therefore yields absorption and flow from one measured field, sharing probe geometry, detector, acquisition clock and photon-arrival axis. It also enables internal short-TOF regression: an Early-gate signal from the same field can enter the Late-gate model as a nuisance regressor. Early-to-late time-window combinations and ratios have been developed for absorption-based depth selectivity [20,21], but their use as a same-field regression basis for TOF-resolved flow remains largely unexplored.

Here we establish how two estimators recovered from the same field retain physiological response amplitude along the same photon-arrival axis (Fig. 1). A controlled forearm cuff perturbation provides the primary within-participant test of the readout-by-TOF interaction. A layered forearm model connects photon time to geometry-conditioned transport weighting. A prefrontal working-memory pilot then extends the same-field analysis to functional activation and introduces short-TOF regression, testing whether the Late-gate BFI response persists after adjustment with an internal Early-gate signal. Together, these experiments turn gate selection into a joint measurement-design problem defined by one detected field, one TOF axis and two fundamentally different estimators. Complete analytical yield across both experiments supports the robustness of the acquisition and analysis workflow, while the network-synchronized seated PFC paradigm demonstrates a route to scanner-free functional measurements in natural postures, complementary to rather than a replacement for fMRI.

**Figure 1.**
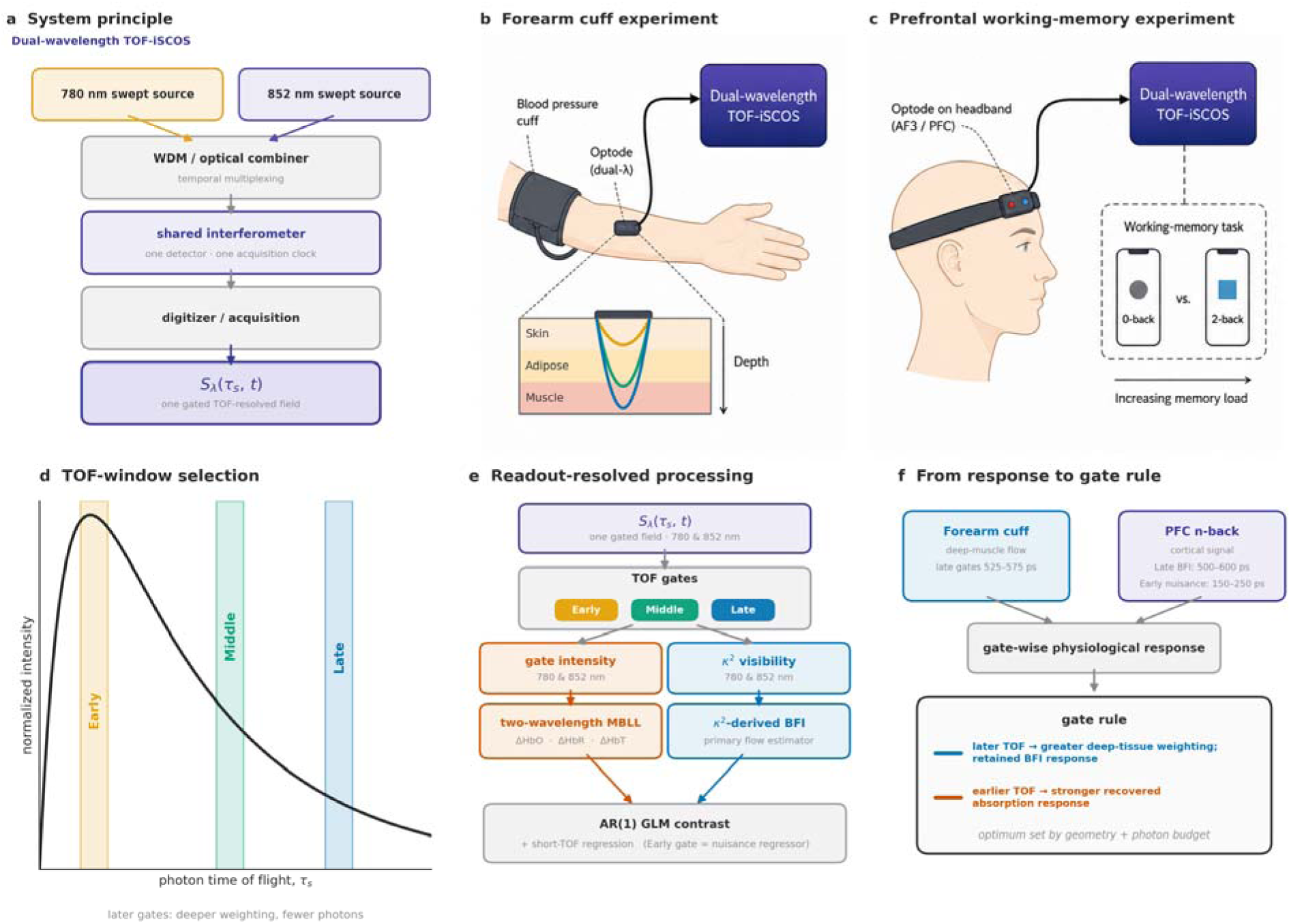
One measurement architecture, two anatomical settings and readout-resolved photon-time analysis. **a.** Dual-wavelength TOF-iSCOS architecture. The 780 and 852 nm swept sources were sequentially multiplexed through a passive fiber WDM combiner into a shared interferometer, detector, digitizer and acquisition clock, yielding a wavelength- and TOF-resolved complex field. **b.** Forearm cuff experiment using a 10-mm source–detector pair to probe a controlled vascular perturbation. **c.** Prefrontal working-memory experiment using the same measurement principle at AF3 during alternating 0-back and 2-back task blocks. **d.** Photon-time selection along the measured TPSF. Early, Middle and Late gates sample overlapping photon-path distributions with progressively greater weighting of longer paths and decreasing photon support toward the TPSF tail. **e.** Readout-resolved processing of the same gated field. Gate-integrated intensity provides the two-wavelength MBLL branch for hemoglobin estimation, whereas temporal field dynamics provide the BFI branch; later-gate task responses can additionally be adjusted using an Early-gate signal through short-TOF regression. **f.** Readout-specific gate design. Controlled forearm measurements establish that absorption and flow retain physiological response differently across photon TOF, while transport modeling shows that gate utility is further conditioned by anatomy, measurement geometry and photon budget. Together, these observations motivate separate photon-time optimization for hemoglobin and flow rather than imposing one common gate on both readouts.

## 2. Results

The interferometric acquisition recovered wavelength-resolved TPSFs together with the complex optical field along the photon time-of-flight (TOF) axis (Fig. 1). Gate-integrated TPSF intensity provided the attenuation signal used for absorption and hemoglobin estimation, whereas repeated complex-field measurements within the same gates provided the temporal statistics used to estimate BFI. Early, intermediate and late gates therefore sampled progressively longer photon paths within the same acquisition while entering distinct intensity- and dynamics-based estimators. This common measurement origin allowed their TOF dependence to be compared without differences in instrumentation or measurement geometry.

### 2.1. Transport modeling defines the depth–photon-support trade-off across photon time-of-flight

Layered Monte Carlo modeling quantified the redistribution of tissue sampling across progressively later model-sampling windows (Fig. 2). The nominal four-layer forearm model comprised 1.5 mm of skin, 3.5 mm of adipose tissue, 15 mm of muscle and an underlying bone layer, with a 5-mm surface-to-muscle depth and 10-mm source–detector separation representing the experimental forearm sampling geometry. From the early to the late model window, the fraction of photon pathlength traversing muscle increased from 2.69% to 21.52%. Across the same windows, muscle-to-superficial dynamic-scattering specificity increased 13.8-fold, compared with a 9.9-fold increase in absorption specificity. This gain in target weighting was accompanied by a 3.4-fold reduction in accepted photon weight. Thus, later photon TOFs simultaneously increased deep-tissue weighting, preferentially increased dynamic-scattering specificity relative to absorption, and reduced photon support, defining a readout-dependent transport trade-off before analysis of the experimental responses.

**Figure 2.**
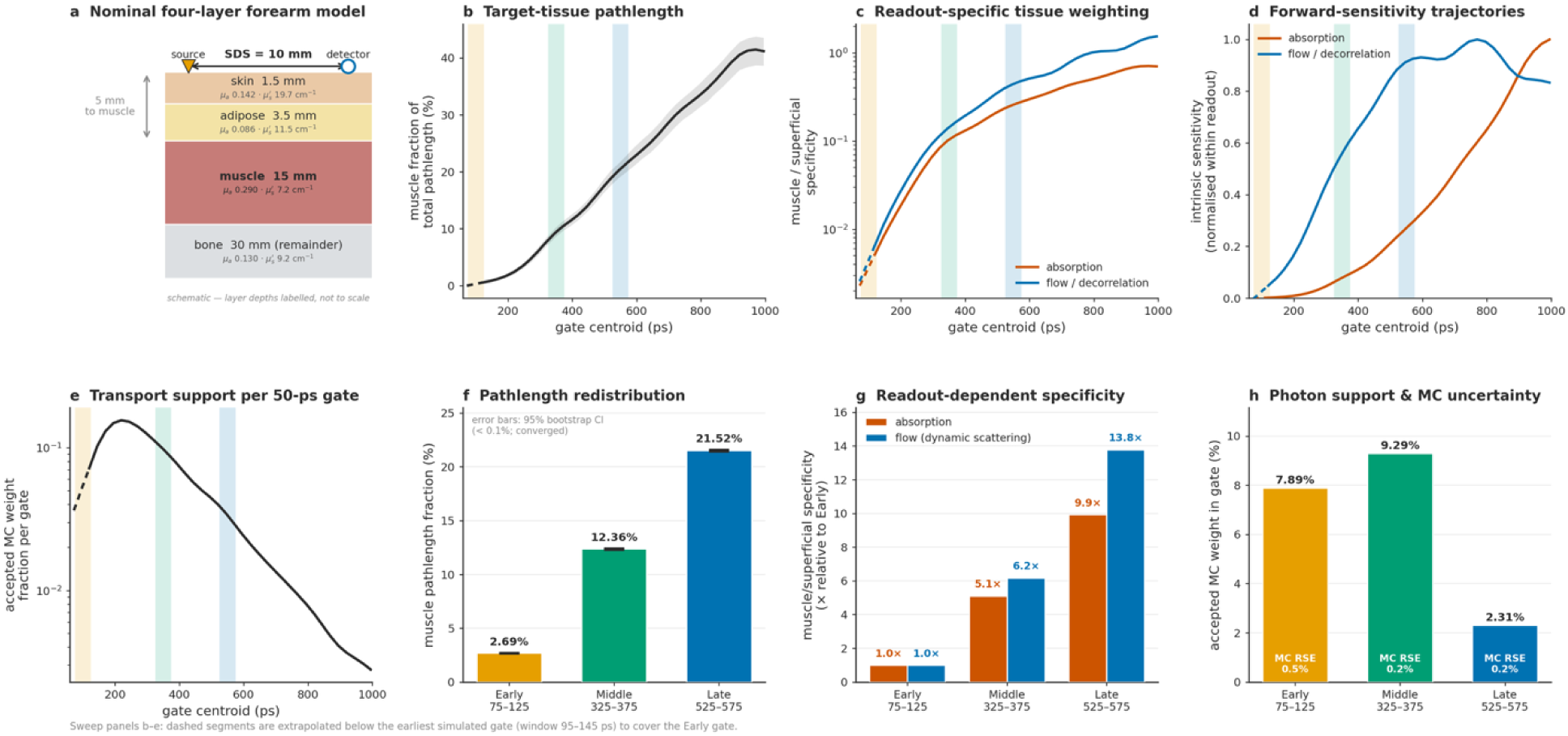
Gate-dependent tissue sensitivity in a nominal four-layer forearm transport model. **a.** Monte Carlo geometry at 780 nm with 5-mm surface-to-muscle depth and 10-mm source–detector separation. **b.** Fraction of detected-photon pathlength accumulated in muscle across photon time-of-flight. **c**, Muscle-to-superficial specificity for absorption and dynamic scattering derived from the same detected photon histories. **d.** Intrinsic target sensitivity normalized within each readout. **e.** Relative Monte Carlo photon support within a 50-ps gate as a function of gate centroid. Shaded bands in **b–e** indicate the experimental Early, Middle and Late TOF windows. **f–h.** Corresponding values evaluated directly within the experimental gates: muscle pathlength fraction (f), muscle-to-superficial specificity normalized to Early (**g**), and relative photon support with Monte Carlo sampling uncertainty (**h**). Simulations used 1.0×10^11^ launched photon histories and the measured instrument response function.

### 2.2. Cuff occlusion separates flow and absorption gate preference

Eleven healthy adults completed a protocol comprising 2.5 min of baseline, 2.0 min of suprasystolic cuff occlusion at 180 mmHg, and post-release monitoring to approximately 7 min total recording time. Cuff inflation produced the expected suppression of BFI together with increased absorption-derived hemoglobin signals, followed by reactive hyperemia after release (Fig. 3). These response directions were preserved across all three 50 ps TOF gates and all 11 participants contributed to the cohort analysis.

**Figure 3.**
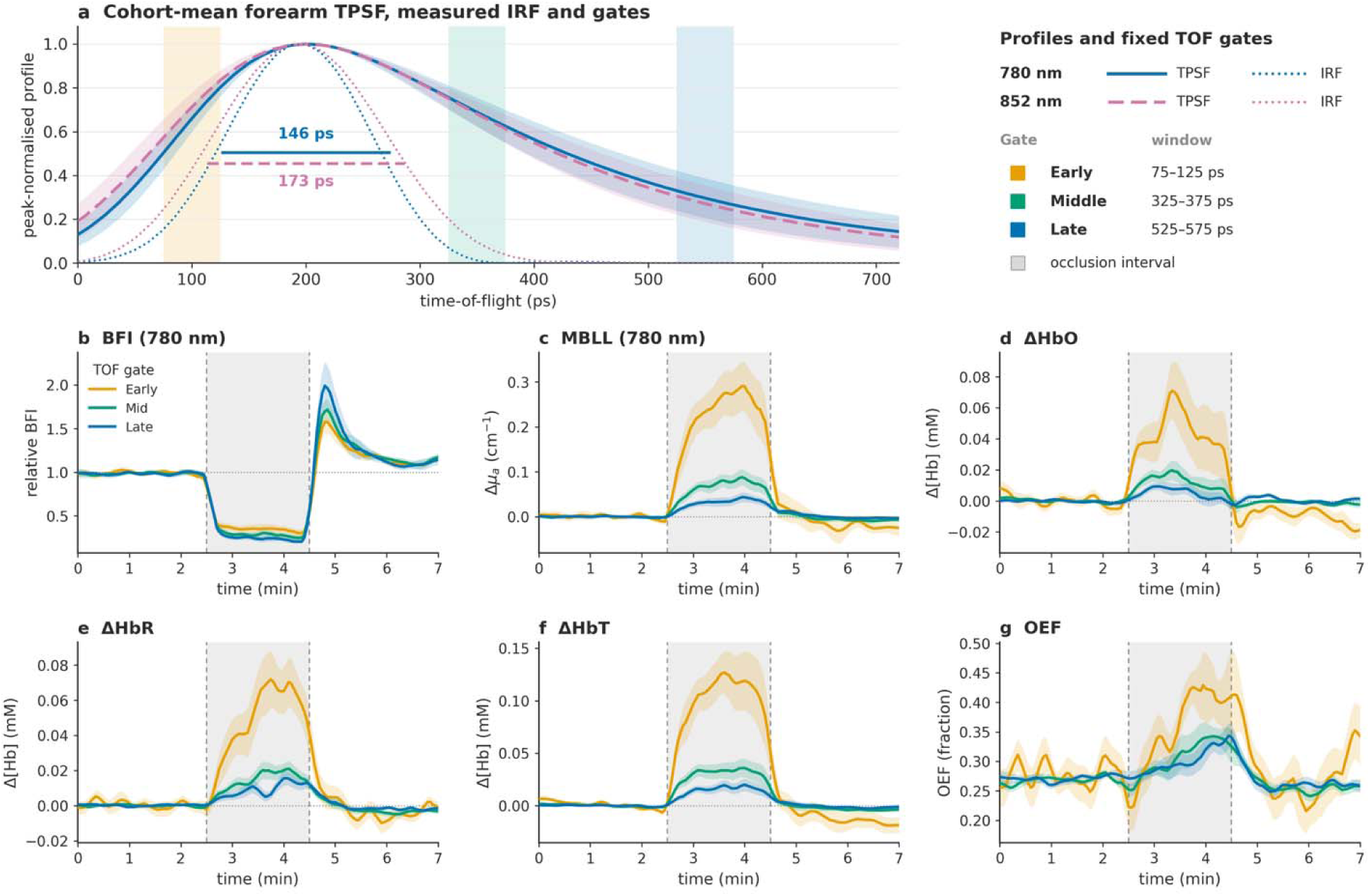
Controlled forearm responses across photon-time gates. **a.** Cohort-mean baseline TPSFs from 11 participants, peak-normalized within participant and wavelength and aligned to 200 ps for display; shading shows ±1 participant SD. Experimental TOF gates and measured IRFs are overlaid. **b-g.** Group trajectories for BFI, Δµ⍰, ΔHbO, ΔHbR, ΔHbT, ΔHbO, ΔHbR, ΔHbT and the descriptive OEF proxy during baseline, suprasystolic cuff occlusion, and release. Lines show participant medians and shaded regions robust standard errors; traces are lightly smoothed for display. The OEF trajectory is an assumption-dependent relative proxy as defined in Methods.

The response amplitudes, however, showed markedly different dependence on photon time-of-flight. BFI suppression during occlusion and the post-release hyperemic response remained well preserved in the Middle and Late gates, whereas absorption and hemoglobin changes were largest in the Early gate and progressively attenuated at later photon TOFs. Thus, although later gates increased modeled muscle weighting for both absorption and dynamic scattering, the experimentally recovered flow and absorption responses retained physiological contrast differently across TOF.

### 2.3. Forearm responses reveal a readout-by-TOF interaction

Participant-level responses confirmed the divergent TOF dependence using the same record-specifi stable-occlusion window and the same mean-response functional for both readouts (Fig. 4). The analysis window comprised the final 1 min before the automatically detected cuff release in each record. Mean BFI suppression at 780 nm increased from 0.631 in the Early gate to 0.698 in the Late gate (two-sided paired Wilcoxon P = 9.8 × 10⁻⁴); at 852 nm, it changed from 0.583 to 0.618 (P = 0.206). Over the identical samples, mean absorption decreased from 0.236 to 0.0397 cm⁻¹ at 780 nm and from 0.218 to 0.0359 cm⁻¹ at 852 nm (P = 0.00195 at both wavelengths). The resulting readout-by-TOF interaction, computed within wavelength and then averaged across wavelengths, was positive in all 11 participants (median 2.61 octaves, 95% bootstrap CI 2.44–3.22; exact two-sided sign-test P = 9.8 × 10⁻⁴). The interaction remained positive in 11/11 participants when the nominal 3.5–4.5 min occlusion window was used (median 2.68 octaves).

**Figure 4.**
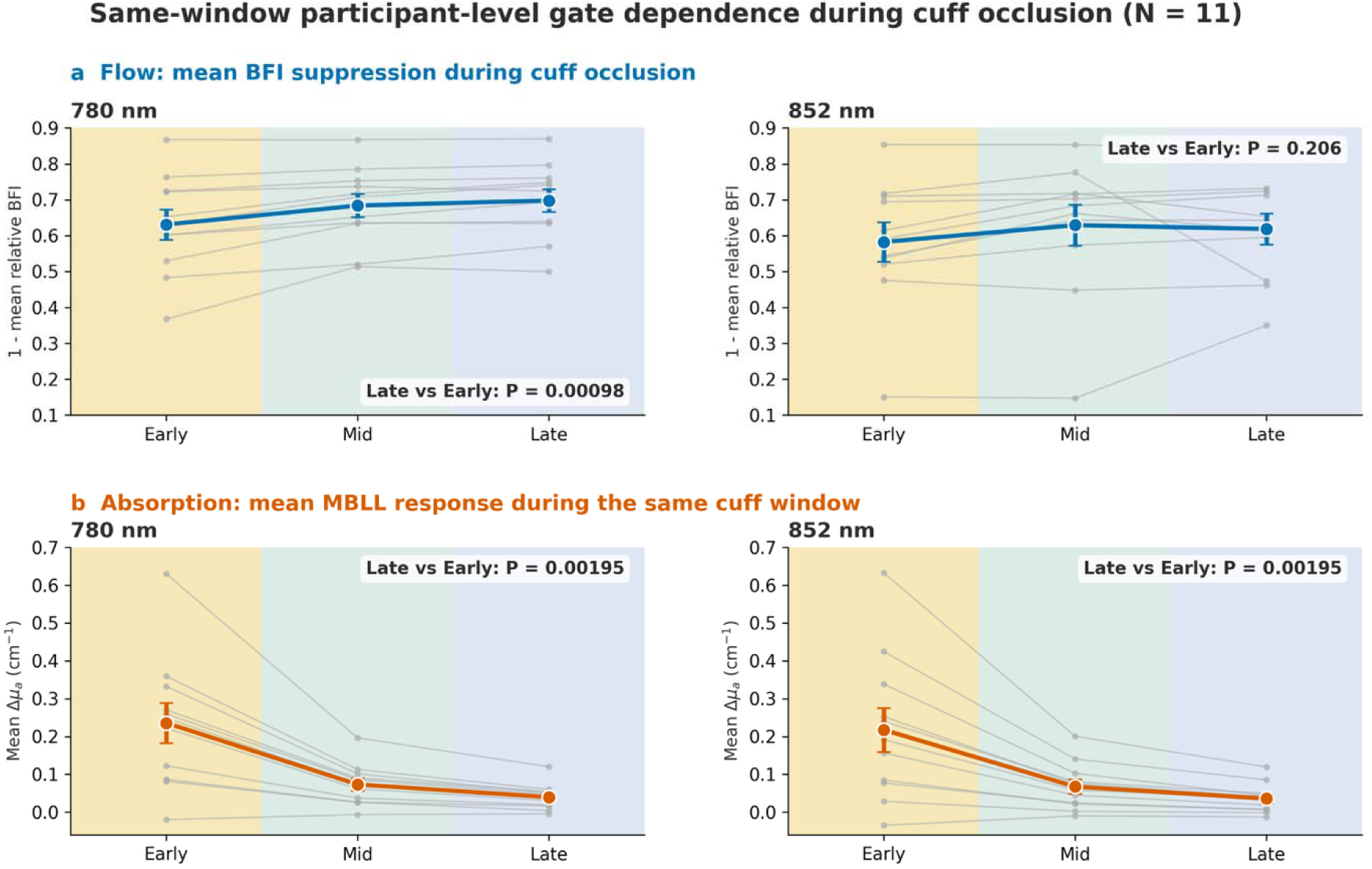
Participant-level readout dependence across photon-time gates during cuff occlusion. Mean BFI suppression (**a**) and MBLL absorption response (**b**) were calculated over identical record-specific stable-occlusion samples in 11 participants. Gray lines connect participant-level responses; colored points show group mean ± standard error. At 780 nm, BFI suppression increased from Early to Late (two-sided paired Wilcoxon P = 9.8 × 10⁻⁴), whereas the corresponding change at 852 nm was not significant (P = 0.206). Absorption decreased from Early to Late at both wavelengths (P = 0.00195). The wavelength-averaged readout-by-TOF interaction was positive in 11/11 participants (median 2.61 octaves, 95% bootstrap CI 2.44–3.22; exact two-sided sign-test P = 9.8 × 10⁻⁴).

### 2.4. Readout-specific gating improves held-out response retention

We next tested whether readout-specific gates retained more physiological response than a single common gate without selecting and evaluating a gate on the same participant. Within each leave-one-participant-out training set, response magnitude was normalized to the largest gate-wise magnitude for each readout and participant. A readout-specific rule selected the gate with the largest training-set median separately for flow and absorption, whereas a common rule selected a single gate after pooling both readouts. Every training fold selected the Late gate for reactive-hyperemia BFI and the Early gate for absorption, whereas the common rule selected the Early gate (Fig. 5). In held-out participants, readout-specific selection increased median pooled normalized response retention from 0.801 to 1.000, corresponding to a median relative gain of 24.8% (participant-bootstrap 95% CI 8.8–42.5%). Thus, the readout-by-TOF interaction translates directly into an engineering penalty when both readouts are forced through one common photon-time window.

**Figure 5.**
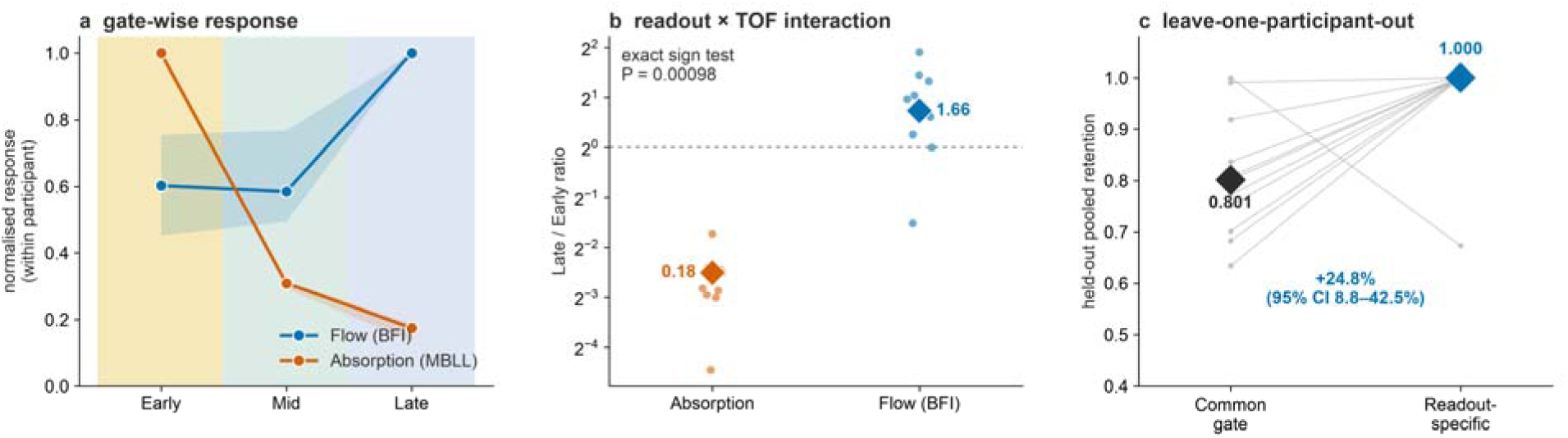
Held-out response retention with common and readout-specific photon-time gates. **a.** Gate-wise response magnitude normalized to the within-participant maximum; markers and error bars show median and interquartile range (N = 11). Flow uses the maximum QC-accepted reactive-hyperemia response within 45 s after release, whereas absorption uses the stable-occlusion mean. **b.** Within-participant Late/Early response ratios for absorption and BFI. **c.** Leave-one-participant-out comparison of a single common gate with readout-specific gate selection. Readout-specific selection increased median pooled held-out response retention from 0.801 to 1.000, corresponding to a 24.8% median relative gain (95% CI 8.8–42.5%).

### 2.5. Prefrontal working-memory responses favor late-gate flow

Seven participants completed a prefrontal 2-back versus 0-back working-memory protocol with the probe positioned over the left prefrontal cortex at AF3 (Fig. 6). The exploratory primary flow endpoint was the unadjusted 2-back minus 0-back BFI contrast from the 500–600 ps TOF window. Two short-TOF sensitivity analyses then tested whether this contrast persisted after accounting for the contemporaneous 150–250 ps Early signal: a joint nuisance-regression model and a complementary rest-calibrated residual analysis. We refer to these Early-to-Late adjustment procedures as short-TOF regression because the nuisance signal is selected along the photon-time axis rather than acquired from a separate source–detector distance.

**Figure 6.**
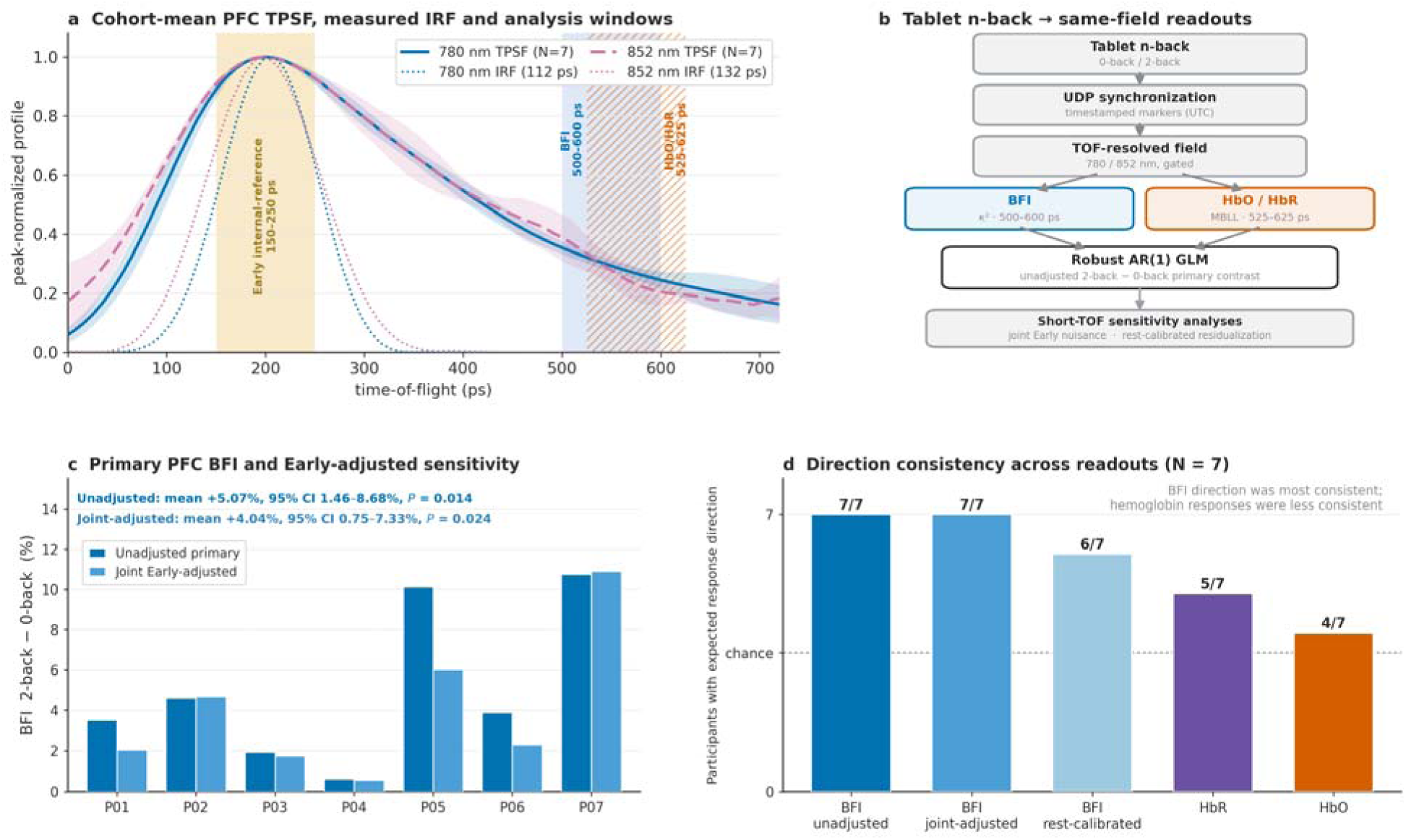
Prefrontal working-memory responses from the same TOF-resolved field. **a.** Cohort-mean baseline PFC TPSFs from seven participants, peak-normalized within participant and wavelength and aligned for display; shading shows ±1 participant SD. Early internal-reference, flow and hemoglobin windows and measured IRFs are overlaid. **b.** Tablet-based 0-back and 2-back task events were synchronized to the acquisition using timestamped UDP markers. The shared TOF-resolved field was processed into BFI and two-wavelength MBLL hemoglobin branches followed by robust AR(1) GLM analysis with short-TOF regression. Separate HRF-convolved regressors represented the 0-back and 2-back task epochs, with intervening rest periods forming the implicit baseline. **c.** Participant-level 2-back minus 0-back BFI contrasts at 500–600 ps. The contrast was positive in 7/7 participants (mean +5.07%, P=0.014) and remained positive in 7/7 after joint Early-gate adjustment (mean +4.04%, P=0.024). **d.** Participant-level expected-direction consistency across readouts. The expected response direction was observed in 7/7 participants for primary BFI, 6/7 after the alternative rest-calibrated BFI adjustment, 5/7 for ΔHbR, and 4/7 for ΔHbO. The dashed line indicates chance-level directional consistency.

Late-gate flow provided the most consistent working-memory response. The 2-back minus 0-back BFI contrast at 500–600 ps was positive in 7/7 participants (mean +5.07%, median +3.89%, 95% t interval 1.46–8.68%; one-sample t-test P = 0.014; exact two-sided sign-test P = 0.0156). The contrast remained positive in 7/7 participants after joint Early-gate nuisance adjustment (+4.04%, 95% t interval 0.75– 7.33%; t-test P = 0.024; sign-test P = 0.0156) and in 6/7 after rest-calibrated residualization (+3.69%, 95% t interval 0.25–7.13%; t-test P = 0.039; sign-test P = 0.125).

Hemoglobin responses were less consistent across participants. In the 525–625 ps gate, ΔHbR was negative in 5/7 participants (mean −2.84 ×10⁻³ mM, P = 0.22), whereas ΔHbO was positive in 4/7 (mean +3.19 ×10⁻³ mM, P = 0.27). Thus, within the same TOF-resolved field measurement, late-gate BFI provided the clearest participant-consistent working-memory contrast in this cohort.

The load manipulation produced the expected behavioral difference, with lower 2-back accuracy and longer reaction times than in 0-back. Within-session adaptation and additional robustness analyses are reported in Supplementary Section S1 and Supplementary Figs. S1–S2.

## 3. Discussion

The central finding is a robust readout-by-TOF interaction within a single interferometric measurement. Using the same participants, stable-occlusion samples, and response functional, Late-gate BFI suppression was retained whereas MBLL absorption progressively declined. The interaction was positive in all 11 participants and remained stable under alternative time-window and preprocessing sensitivity analyses. Layered transport modeling showed that later photon TOFs progressively increased muscle weighting and preferentially increased dynamic-scattering specificity relative to absorption, while reducing photon support. The experimental divergence therefore reflects both readout-dependent forward sensitivity and readout-dependent estimator behavior within the same photon-limited measurement. Because tissue thicknesses were nominal rather than participant-specific, the simulated fractions should be interpreted as geometry-conditioned sensitivity estimates rather than subject-specific anatomical measurements. Photon time-of-flight is consequently not only a depth coordinate, but a readout-dependent measurement coordinate whose utility depends on the estimator applied to the detected field.

This distinction is particularly relevant for hybrid systems that recover hemoglobin and flow from the same acquisition. MBLL remains attractive because it converts wavelength-resolved intensity changes into familiar hemoglobin contrasts using a simple and computationally efficient estimator and underpins conventional continuous-wave fNIRS [5,8,9]. In the present forearm measurements, however, the largest recovered absorption and hemoglobin responses occurred in the Early gate and decreased strongly with photon TOF, reaching a median absorption Late/Early ratio of 0.18. BFI showed a markedly different dependence: physiologically expected occlusion and reactive-hyperemia responses remained measurable in later gates despite lower photon support. Previous TD-DCS studies have similarly shown that flow-sensitive gate selection involves a balance between increasing late-photon tissue sensitivity and declining photon statistics and instrument-response constraints [22,24,26,27]. This does not imply that BFI is universally more noise-efficient than MBLL. Rather, it demonstrates that increased target weighting at later photon TOFs does not translate into equivalent retained response amplitude for different estimators. Notably, the Early-to-Late increase in cuff BFI was significant at 780 nm but not at 852 nm, where late-gate photon support was lower. The wavelength-averaged interaction nevertheless remained positive in all participants, reinforcing that gate utility is conditioned by photon budget and estimator performance rather than by TOF alone.

The engineering consequence was confirmed by the held-out gate-selection analysis. When a single gate was selected after pooling the two readouts, the Early gate was preferred because of the strong absorption response. When gates were selected independently, every training fold selected Early for absorption and Late for reactive-hyperemia BFI. Readout-specific selection increased held-out pooled response retention from 0.801 to 1.000, corresponding to a median gain of 24.8%. Thus, forcing hemoglobin and flow through one photon-time window imposes a measurable performance penalty even though both originate from the same detected field. A practical hybrid architecture should therefore retain the simplicity of MBLL while allowing a separately optimized flow branch.

The prefrontal working-memory measurements provide an initial functional extension of this principle. The 500–600-ps BFI contrast between 2-back and 0-back was positive in all seven complete sessions and remained positive in all seven after adjustment using the contemporaneous Early-TOF signal. Hemoglobin responses were less consistent across participants. Flow and hemoglobin are known to provide physiologically distinct vascular contrasts, with perfusion-sensitive and hemoglobin-based signals differing in temporal and vascular-compartment weighting [5,8,9], conceptually analogous to distinctions between perfusion- and BOLD-weighted functional MRI [36–38]. The Early-to-Late adjustment used here, which we term short-TOF regression, differs conceptually from conventional short-separation regression: nuisance and target signals are obtained from the same source–detector geometry and detection channel but from different regions of the photon-time distribution. The Early signal is therefore an overlapping-path internal proxy rather than an independent superficial measurement. Persistence of the Late-gate BFI contrast after this adjustment demonstrates robustness to shared Early-gate covariance, but does not by itself establish cortical localization or superiority over an independent short-separation channel.

These findings complement previous work showing that time-domain and correlation-based flow measurements balance late-photon tissue sensitivity against photon statistics and instrument response [22,24,26,27], and that NIRS and DCS can exhibit modality-dependent tissue sensitivity [19]. Functional TD-DCS, massively parallel DCS, interferometric diffusing-wave spectroscopy, and high-density SCOS have demonstrated functional flow responses using dedicated flow architectures, wider gates, longer separations, multiple detectors, or superficial reference channels [39–42]. CoMind R1 recently demonstrated TOF-dependent visual-cortex hyperemia in 16 adults using relative BFI as the principal functional endpoint [43]. The present study addresses a complementary question by placing hemoglobin and flow estimators on the same measured TOF axis. Dual-wavelength absorption and hemoglobin are recovered together with TOF-resolved BFI from the same interferometrically detected field, enabling readout-specific gate design and same-field short-TOF adjustment within one acquisition architecture.

The demonstrated effect is readout dependence rather than a universal gate boundary in picoseconds. Absolute gate choice will depend on anatomy, source–detector geometry, IRF, photon budget, wavelength, and estimator conditioning. The present study used a nominal forearm geometry and a single AF3 prefrontal location, relative flow quantification, temporally interleaved wavelengths, and no IRF deconvolution; tissue thickness was not measured individually. The PFC cohort was small and establishes response consistency rather than spatial localization or population-level generality. Future studies should combine anatomy-conditioned transport modeling with improved photon collection, systemic monitoring, independent short-separation measurements, and multi-channel cerebral coverage in larger cohorts. Within these boundaries, the current results establish a broader design principle: photon time-of-flight should be optimized for the readout being recovered rather than imposed as a single common window across physiologically distinct observables.

## 4. Methods

### 4.1. Dual-wavelength interferometric measurement

Two fiber-coupled distributed-feedback lasers were used: 780 nm (EYP-DFB-0780-00020-1500-BFY12-0005, Eagleyard–Toptica) and 852 nm (EYP-DFB-0852-00015-1500-BFY12-0005, Eagleyard–Toptica). Their outputs were combined with a three-wavelength fiber WDM (RNN50HA, Thorlabs; 642 nm port unused) and routed through a shared Mach–Zehnder measurement interferometer. Diffusely backscattered light was collected into a single-mode fiber, recombined with the reference field and detected with a balanced detector; a separate fixed-path reference interferometer supplied the phase signal for optical-frequency linearization. Delivered powers were 11 mW at 780 nm and 9 mW at 852 nm, approximately 2.5-fold and 4.3-fold below the respective ANSI Z136.1-2014 maximum permissible exposures. The forearm source–detector separation was 10 mm. Cohort-mean baseline TPSFs were computed after within-participant, within-wavelength peak normalization and peak alignment to 200 ps for display; shaded bands show ±1 participant SD. The fixed gates and acquisition-derived wavelength-specific IRF profiles are shown with the corresponding forearm and prefrontal results in Figs. 3 and 6.

The wavelengths were acquired in sequential, prospectively labeled chunks; only one source was swept at a time, so the channels were temporally interleaved rather than simultaneously mixed and later demultiplexed. Signals were digitized synchronously with a Spectrum M2p.5962-x4 16-bit digitizer at 100 MS s⁻¹ per channel and a 50 kHz sweep rate. The first 150 ms after each wavelength transition were discarded. The mean same-wavelength revisit interval was 2.94 s (effective wavelength-specific sampling rate about 0.34 Hz), and all estimates retained acquisition timestamps. Each wavelength was independently linearized, resampled and Fourier transformed to recover the complex TOF-resolved field. Calibration was acquisition-specific. For the representative forearm record, the raw TOF-bin spacings were 12.61 ps at 780 nm and 14.96 ps at 852 nm, and the averaged acquisition-stored IRF profiles had FWHMs of 146.0 ps and 173.1 ps, respectively. For the representative PFC session, the corresponding raw spacings were 9.68 ps and 11.46 ps and the IRF FWHMs were 112.1 ps and 132.4 ps; the wavelength-specific PFC outputs were subsequently registered to the common corrected photon-time axis used for gating. No IRF deconvolution was applied [28–30].

### 4.2. Photon-time gates

The forearm analysis used three development-selected 50-ps windows on the registered zero-TOF axis: Early, 75–125 ps; Middle, 325–375 ps; and Late, 525–575 ps. The 550-ps Late center was selected during estimator development with consideration of late-gate 852-nm fit yield, then frozen before full-cohort inference; the analysis was not externally preregistered. The prefrontal analysis used an Early 150–250-ps internal nuisance window, a 500–600-ps flow window and a 525–625-ps hemoglobin window. Absolute picosecond values are instrument- and geometry-specific and are not treated as universal depth strata.

### 4.3. Monte Carlo transport model

GPU-accelerated Monte Carlo photon transport was performed using MCX [45] in a nominal four-layer forearm geometry at 780 nm. The model comprised 1.5 mm of skin, 3.5 mm of adipose tissue, 15 mm of muscle and a 30 mm deep bone-proxy layer in a 120 × 120 × 50 mm volume sampled with 0.5 mm isotropic voxels. Layer-specific absorption and reduced-scattering coefficients are listed in Table 1; g=0.90 and n=1.40 were used for all tissues. MCX scattering coefficients were obtained as 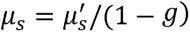. The source was a normally incident pencil beam, and the detector was 1 mm in diameter with NA = 0.39 at a source–detector separation of 10 mm. Refractive-index mismatch was enabled. A total of 1.0 × 10^11^ photon histories were launched.

**Table 1.** Layer-specific properties used in the 780-nm forearm Monte Carlo model. An anisotropy factor of g=0.90 and refractive index of n=1.40 were used for all layers. The bone layer extended to the lower boundary of the simulation domain. Skin, adipose, and muscle optical properties were derived from Sayli et al. [48]; bone was treated as a fixed boundary-tissue proxy. Dynamic parameters were literature-derived population priors rather than participant-specific estimates.

| Layer | Thickness (mm) | $\mu_a \text{ (cm}^{-1}\text{)}$ | $\mu'_s \text{ (cm}^{-1}\text{)}$ | $\alpha D_b \text{ (cm}^2\text{/s)}$ |
| --- | --- | --- | --- | --- |
| Skin | 1.5 | 0.1422 | 19.7420 | $1.0 \times 10^{-8}$ |
| Adipose | 3.5 | 0.0862 | 11.4720 | $5.0 \times 10^{-9}$ |
| Muscle | 15 | 0.2896 | 7.1660 | $5.0 \times 10^{-8}$ |
| Bone | 30 | 0.1300 | 9.1538 | $2.0 \times 10^{-10}$ |

Monte Carlo photon transport was simulated at nominal optical properties, and the same detected photon histories were subsequently used for independent absorption- and dynamic-scattering sensitivity analyses. For each detected photon, MCX retained time of flight, photon weight, tissue-resolved pathlength and tissue-resolved momentum transfer. Before gate assignment, photon arrival times were convolved with the acquisition-stored 780-nm forearm IRF from the cohort-medoid P08-L recording (mean of 156 stored IRFs; FWHM, 146.0 ps), after edge-median baseline correction, clipping to non-negative values, peak alignment and normalization to unit area. Continuous TOF trajectories were evaluated using a sliding 50-ps window on the registered photon-time axis, and gate-wise summary metrics were calculated directly within the experimental Early (75–125 ps), Middle (325–375 ps) and Late (525–575 ps) windows. The displayed extension below the earliest directly simulated sweep window was included only for visual continuity and was excluded from all gate-wise calculations. Gate-wise photon support was defined as the fraction of total accepted Monte Carlo weight assigned to each window. Monte Carlo precision was quantified using the larger of the relative standard errors of the mean muscle pathlength and the muscle pathlength fraction; the relative standard error did not exceed 0.5% in any of the three experimental gates.

Absorption sensitivity was evaluated by post hoc reweighting of the detected histories without altering photon trajectories. For a perturbation Δ*μ_a,k_* in tissue *k* photon weight was updated as

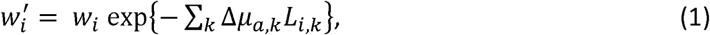

Where *L_i,k_* is the pathlength of photon *i* within tissue *k*. Nominal absorption was evaluated together with global ±10% scaling of the tissue-specific baseline *μ_a_* values. Muscle absorption sensitivity within each TOF gate was quantified from the IRF- and gate-weighted muscle pathlength,

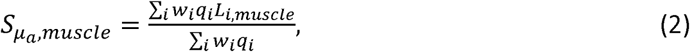

Where q*_i_* denotes the probabilistic contribution of photon *i* to the selected gate after IRF convolution.

Dynamic-scattering sensitivity was evaluated from the same photon histories using a DCS-like correlation model in which flow was represented by the effective dynamic-scattering parameter *αD_b_*, rather than by explicit fluid dynamics. Baseline *αD_b_* values were 1×10^-8^, 5×10^-9^, 5×10^-8^ and 2×10^-10^ *cm*^2^*s*^-1^ for skin, adipose tissue, muscle and bone, respectively. For photon *i*, the decorrelation rate was

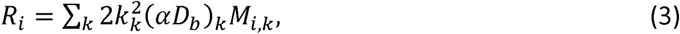

Where *M_i,k_* is the accumulated momentum transfer in tissue *k* and *k_k_* = 2*πn_k_*/*λ*. The gate-resolved field autocorrelation was then

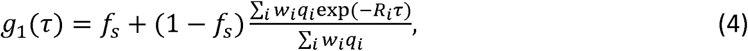

With *f_s_* = 0.05.

Dynamic-scattering sensitivity was estimated numerically using 10% perturbations applied separately to muscle, to the superficial compartment (skin and adipose tissue), and to both compartments. Muscle sensitivity was defined as

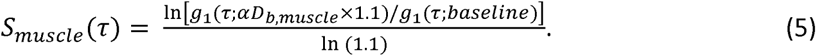

Because increasing *αD_b_* accelerates field decorrelation, this sensitivity is negative. Its magnitude was therefore summarized across correlation delay using the RMS value, and dynamic-scattering specificity was defined as

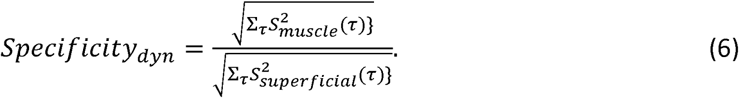

Absorption and dynamic-scattering analyses therefore interrogate different physical properties of the same detected photon ensemble: absorption sensitivity reflects tissue-resolved photon pathlength, whereas dynamic-scattering sensitivity reflects the tissue-resolved momentum-transfer events that contribute to field decorrelation. The simulations quantify intrinsic gate-dependent sensitivity and specificity and are not intended as a direct forward simulation of cuff-response amplitude.

### 4.4. Subjects

All participants were healthy adults with no known neurological or cardiovascular disease and were recruited at a single center (Table 2). Eleven adults contributed 12 uninterrupted forearm recordings; one participant was measured bilaterally, and the two records were averaged before participant-level inference. Seven participants contributed seven complete prefrontal cortex (PFC) sessions, with one analyzed session per participant. No complete session was repeated or excluded based on the optical response. The 500–600 ps PFC flow gate was fixed after the first five complete sessions and applied unchanged to the final two-session extension. Participants provided written informed consent. All procedures were approved by the Commission of Bioethics at the Military Institute of Medicine, Poland (permission no. 90/WIM/2018) and were conducted in accordance with the Declaration of Helsinki. Participant and protocol reporting followed fNIRS best-practice recommendations where applicable [44].

**Table 2.** Participant characteristics (group summaries). Forearm summaries use the 11 independent participants retained in the present analysis. All prefrontal participants were male. Bilateral forearm recordings from one participant were averaged before participant-level inference.

| Characteristic | Forearm cohort | Prefrontal cohort |
| --- | --- | --- |
| Independent participants | 11 (12 recordings; bilateral averaged) | 7 participants (7 complete sessions) |
| Sex | 9 male, 2 female | 7 male |
| Age | 36.6 ± 6.8 years; median 40; range 24–44 | 24–42 years (n = 7) |
| BMI | 25.3 ± 3.3 kg m<sup>-2</sup>; median 24.2; range 21.0–32.0 | 23.7 ± 1.6; median 24.0; range 21.0–24.9 |
| Fitzpatrick skin type | median II; range II–V (II n = 8, III n = 1, IV n = 1, V n = 1) | range II–IV (II, n = 6; IV, n = 1) |

### 4.5. Forearm cuff protocol

Participants were seated with the measured forearm supported. The optical probe was positioned on the ventral forearm at a 10 mm source–detector separation, and a pneumatic cuff was placed on the ipsilateral upper arm above the elbow. After 2.5 min of baseline, the cuff was inflated to 180 mmHg for 2.0 min and then released; recording continued to approximately 7.5 min. Baseline normalization used the 0.5–2.0 min interval.

Cuff release was identified automatically from the maximum positive temporal derivative of the smoothed median 780 nm BFI across TOF gates within a predefined ±0.75-min interval around the nominal 4.5 min release time. The primary stable-occlusion endpoint was calculated over the final 1 min before the detected release; the fixed 3.5–4.5 min interval was retained as a clock-time sensitivity analysis. Reactive hyperemia was quantified as the maximum QC-accepted BFI response within 45 s after release.

### 4.6. Prefrontal working-memory protocol

The optical probe was positioned over the left prefrontal cortex at AF3 of the international 10–10 EEG system. AF3 was localized individually using an EasyCap electrode cap immediately before optical recording and the position was marked on the forehead. Participants performed a tablet-based n-back working-memory task comprising eight 2-back and eight 0-back blocks of approximately 30 s presented in alternating order. The approximately 30-s task blocks were separated by nominal 20-s rest periods. Marker-defined rest samples were not assigned to either load condition and served as the implicit baseline in the session-level model. The 0-back condition served as a low-working-memory-load control, whereas the 2-back condition required comparison of each stimulus with that presented two trials earlier [31–35]. Functional responses were defined from the within-participant 2-back minus 0-back contrast.

Accuracy and reaction time were recorded together with the optical acquisition. Per-block accuracy, hit and false-alarm rates, and median reaction time on detected targets were used to verify the load manipulation and to assess descriptive within-session adaptation. The first four and final four blocks of each load defined the two halves used for the adaptation analysis. Two sessions began with 0-back and five with 2-back.

### 4.7. Stimulus-acquisition synchronization

The tablet n-back application and desktop acquisition were synchronized over the local network using timestamped event markers for session start and end, block, trial, stimulus type and participant response. Both the tablet-device timestamp and the desktop-receipt timestamp were retained to align behavioral events with the TOF-resolved optical acquisition. The full message protocol and event manifest are provided in Supplementary Section S2.

### 4.8. Flow, absorption and hemoglobin estimation

TOF-resolved relative blood-flow index (BFI) was estimated from the temporal dynamics of the reconstructed complex optical field using the finite-exposure correlation analysis described previously [29,30]. BFI was estimated independently within each wavelength and TOF gate and normalized to the corresponding within-recording baseline. For the forearm analysis, frames were retained at R^2^ ≥ 0.80, isolated single-frame gaps were linearly interpolated, and the resulting trajectories were processed with a three-frame moving median followed by a three-frame moving mean. Response windows required at least 70% accepted frames. For the PFC analysis, the processing settings were frozen with the cognitive analysis: fitted BFI values were retained at R^2^ ≥ 0.10, log-domain outliers exceeding five median absolute deviations were removed, isolated single-frame gaps were interpolated, and a three-frame moving median was applied. No additional moving-mean filter or minimum accepted-frame fraction was imposed on the PFC endpoint. Gate- and wavelength-specific fit yield and quality-control summaries are reported in the Supplementary Information.

For absorption analysis, the median TPSF signal at corrected TOF < 0 ps was subtracted from each frame as a pre-arrival background, and negative samples were set to zero. Gate intensity I_G_(t) was calculated by integrating the background-corrected TPSF within gate G. The baseline reference I_0,G_ and reference *TPSF_0_*(*τ_s_*), were calculated over the 0.5–2.0 min baseline interval, with I_0,G_ defined as the gate-integrated baseline TPSF. A gate-specific mean photon time-of-flight and corresponding mean pathlength were calculated from the centroid of the reference TPSF within the gate,

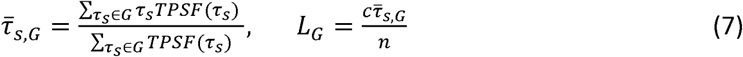

where *τ_s_* denotes photon time-of-flight, c=0.299792458 mm/ps, and n=1.4. Napierian optical-density and absorption changes were then calculated as

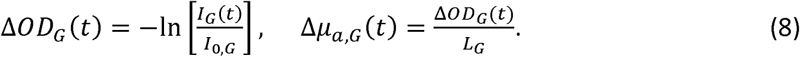

The primary absorption trajectories were processed with a three-frame moving mean; unsmoothed Δ*μ_a_* was retained for sensitivity analysis.

Because the 780 and 852 nm measurements were acquired sequentially rather than simultaneously, the wavelength-specific Δμa traces were registered using their acquisition timestamps before spectroscopic inversion. The 852 nm trace was linearly interpolated onto the 780 nm time grid, and the two-wavelength inversion was restricted to time points with support from both wavelength streams; no extrapolation beyond their common temporal support was used.

Paired 780 and 852 nm absorption changes were converted to relative oxy- and deoxyhemoglobin concentration changes, ΔHbO and ΔHbR, using a two-wavelength modified Beer–Lambert inversion [46,47]. Prahl/OMLC decadic extinction coefficients were converted to Napierian units by multiplication by ln(10) [49]. The tabulated 850-nm coefficients were used for the 852 nm channel as a 2 nm approximation. Hemoglobin concentrations are reported in mM relative to the within-recording baseline, with ΔHbT=ΔHbO+ΔHbR.

For the descriptive forearm OEF proxy only, literature-assigned resting values StO_2,0_=0.73 and HbT_0_=0.119 mM were used, based on resting forearm-muscle measurements reported by Re et al. [50], with HbO_0_=StO_2,0_HbT_0_:

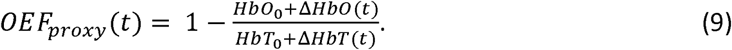

This quantity is an assumption-dependent relative trajectory and is not interpreted as an absolute participant-specific oxygen-extraction measurement.

### 4.9. Short-TOF regression and PFC analysis

For each readout *b* and TOF gate *G*, the 2-back minus 0-back contrast was estimated at the session level using a robust general linear model with AR(1) prewhitening, separate canonical-HRF-convolved regressors for the 0-back and 2-back conditions, and cubic session-time drift terms. Short-TOF regression denotes the inclusion of the contemporaneous Early-gate signal from the same acquisition as an internal nuisance regressor for a later-gate response. The joint short-TOF model was

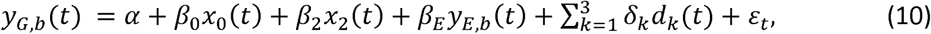

with

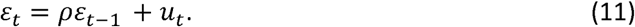

Here, *y_G,b_*(*t*), is the signal for TOF gate *G* and readout *b*, and *α* is the session intercept. The regressors *x*_0_(*t*) and *x*_2_(*t*) were obtained by convolving separate boxcar indicator functions for the marker-defined 0-back and 2-back epochs with the canonical hemodynamic response function. The unconvolved indicators were zero during intervening rest periods. After convolution, the regressors retained the expected hemodynamic carryover into the early portion of each rest period. Rest therefore constituted the implicit reference condition represented by the intercept and drift terms; it was not pooled with either task condition.

The contemporaneous Early-gate signal *y_E,b_*(*t*) was included as an internal nuisance regressor, with coefficient *β_E_*. The coefficients *β*_0_ and *β*_2_ quantify the fitted 0-back and 2-back responses relative to the implicit rest reference. The reported within-session load contrast was *C*_2-0_ = *β*_2_ – *β*_0_.

The functions *d_k_* describe slow within-session drift. Session time was first scaled to

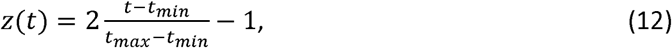

and the polynomial drift basis was defined as 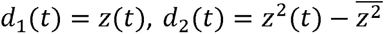, and 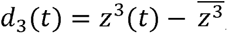, where the overbar denotes the temporal mean over the session. Accordingly, *δ*_1_, *δ*_2_, and *δ*_3_ are nuisance coefficients for the linear, quadratic, and cubic components of session drift, respectively. In Eq. (11), *ρ* is the first-order autoregressive coefficient and *u_t_* is the innovation term. Model coefficients were estimated after AR(1) prewhitening with Tukey-bisquare robust reweighting.

A complementary rest-calibrated analysis estimated the Early-to-Late coupling using samples outside the task blocks,

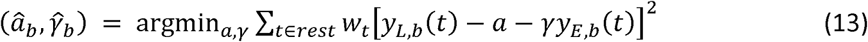

where *L* denotes the Late gate and *W_t_* is the robust-regression weight. The corrected Late-gate signal was then formed as

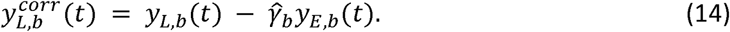

Only the fitted Early-gate contribution was subtracted; the intercept 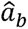 was retained to preserve the Late-gate signal offset. The unadjusted 500–600 ps BFI contrast, estimated from the same session model after omitting the Early-gate nuisance term, was the exploratory primary PFC endpoint. The joint model in Eqs. (10) and (11) was the primary short-TOF sensitivity analysis, whereas the rest-calibrated residualization in Eqs. (13) and (14) provided a complementary sensitivity analysis.

Because the Early and Late TOF windows contain overlapping photon-path distributions, short-TOF regression is not equivalent to conventional short-separation regression using an independent source– detector channel. The PFC BFI estimator and 500–600 ps flow gate were fixed after the first five complete sessions and applied unchanged to the final two-session extension. Development (N=5) and frozen-extension (N=2) results were examined separately as a robustness check.

### 4.10. Statistical analysis

The participant was the biological unit of analysis. Bilateral forearm recordings from one participant were averaged before participant-level inference, and the two wavelengths were not treated as independent replicates.

Paired Early–Late comparisons were evaluated using two-sided Wilcoxon signed-rank tests. The primary forearm endpoint was the readout-by-TOF interaction calculated from identical stable-occlusion samples for flow and absorption. Flow response was defined as

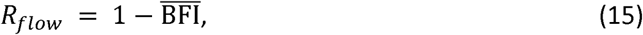

and absorption response as

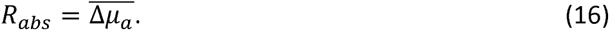

Within each wavelength, the interaction was

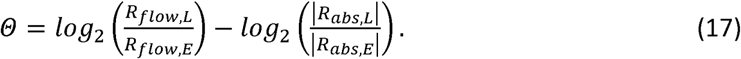

The interaction was then averaged across wavelengths within participant. Its P value was obtained using an exact two-sided sign test, and the 95% confidence interval for the group median was estimated using a 20,000-resample participant bootstrap with a fixed random seed. Sensitivity analyses repeated the interaction using the nominal 3.5–4.5 min occlusion interval and pre-censoring/pre-smoothing trajectories.

The held-out gate-selection analysis used wavelength-averaged response magnitudes: the maximum quality-control-accepted κ^2^-based BFI response within 45 s after cuff release for flow and the absolute stable-occlusion mean Δμ_a_ for absorption. For each participant and readout, the Early, Middle and Late responses were divided by the largest response across the three gates. In each leave-one-participant-out fold, a readout-specific rule selected the gate with the largest training-participant median separately for flow and absorption, whereas a common rule selected one gate by maximizing the median after pooling the two readouts with equal weighting. The selected gates were then applied to the held-out participant, and the held-out score was the median of that participant’s normalized flow and absorption retentions. Relative gain was calculated as (readout-specific retention/common-gate retention) − 1. The reported group values are participant medians; the 95% confidence interval for the median relative gain used 20,000 participant-level bootstrap resamples with a fixed random seed. This secondary analysis quantified normalized response retention rather than absolute detectability or practical gate utility.

PFC contrasts were summarized by participant-level effect estimates and 95% confidence intervals. Two-sided one-sample t-tests were used for exploratory group-level inference, with exact two-sided sign tests used as distribution-light checks of response-direction consistency. The forearm readout-by-TOF interaction was the single primary statistical endpoint. PFC, individual-wavelength, gate-decomposition and hemoglobin analyses were exploratory, and no multiplicity adjustment was applied.

## 5. Conclusions

Dual-wavelength TOF-iSCOS revealed a reproducible readout-by-TOF interaction: under the tested forearm geometry, Late gates retained flow contrast whereas MBLL absorption was strongest in Early gates, and readout-specific gate selection improved held-out response retention. Exploratory prefrontal measurements extended this principle to functional monitoring, with a consistent Late-gate BFI contrast that survived same-field Early-gate adjustment, without establishing cortical localization. Photon time-of-flight should therefore be treated as a readout-dependent measurement coordinate. Hybrid interferometric systems should preserve separate, geometry-conditioned photon-time branches for intensity-based hemoglobin and dynamics-based flow rather than impose a common gate.

## Data availability

Data underlying the results presented in this paper are not publicly available at this time but may be obtained from the authors upon reasonable request.

## Acknowledgements

The authors thank InCellVu for in-kind support through access to laboratory space and the loan of computing hardware for data analysis and optical instrumentation for system calibration and characterization. InCellVu had no role in the study design, data analysis or interpretation, or preparation of the manuscript.

## Author contributions

M.M.: investigation, visualization and writing – original draft. N.M.: investigation, data curation and writing – review and editing. K.N.-P.: methodology, validation and writing – review and editing. D.B.: conceptualization, methodology, supervision, software, optical system, funding acquisition, project administration and writing – original draft, review and editing. All authors reviewed and approved the manuscript.

## Supplementary information

### S1. Behavioral performance and within-session adaptation

The prefrontal cohort (N=7) performed a block-alternating 2-back/0-back protocol with eight blocks per load. The working-memory manipulation was effective: 0-back accuracy was at or near ceiling, whereas 2-back accuracy was 2–10 percentage points lower (participant means, 89.6–95.8%). The reaction-time cost for detected targets, calculated from median affirmative-response times, ranged from approximately +91 to +329 ms (Fig. S1a,b).

**Figure S1.**
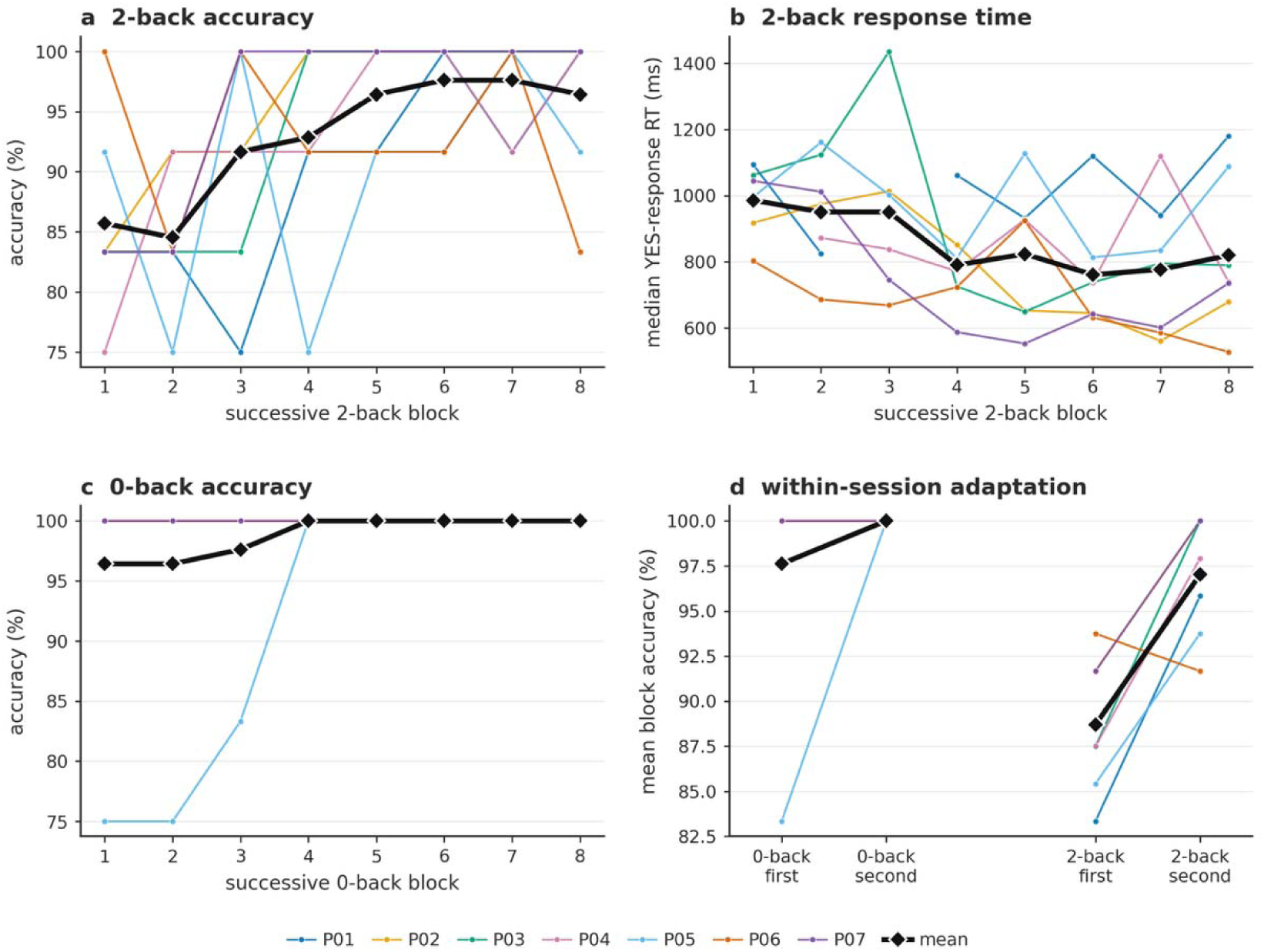
Within-session behavioral adaptation in the prefrontal cohort (N=7). **a.** 2-back accuracy across the eight successive 2-back blocks for each participant (P01–P07) and the cohort mean (black diamonds). **b.** Median response time for affirmative, detected-target responses across successive 2-back blocks; missing points indicate blocks without an affirmative response. **c.** 0-back accuracy across the eight successive 0-back blocks. **d.** Within-participant mean block accuracy in the first and second halves of the session for each load. The largest improvement occurred over the early blocks, consistent with within-session task familiarization.

Performance also showed a clear within-session familiarization effect. Mean 2-back accuracy increased from 88.7% in the first half to 97.0% in the second, corresponding to a mean paired change of +8.33 percentage points. Six of seven participants improved by 8.3–12.5 percentage points, whereas one participant decreased by 2.1 percentage points. Reaction times to detected targets shortened over the early blocks before stabilizing (Supplementary Fig. S1a,b,d). This practice effect motivates a longer familiarization period and explicit counterbalancing of block order in future confirmatory studies.

Across the same session halves, the primary 500–600-ps BFI contrast between 2-back and 0-back decreased from a cohort mean of 9.13% to 3.89% (mean paired change, −5.25 percentage points; median paired change, −3.82 percentage points) and was lower in the second half in five of seven participants (Fig. S2). Thus, group-average behavioral performance improved while the late-gate flow contrast decreased as the task became more familiar. The participant-level directions were heterogeneous, however, and this descriptive N=7 comparison does not establish behavioral–flow coupling or causality.

One protocol deviation occurred. During a single 2-back block, participant P05 reported a phone/tablet notification, around which three behavioral errors clustered. In accordance with the prespecified analysis policy, the block was retained in the primary analysis. Excluding it increased that participant’s 2- back accuracy from 89.6% to 91.7% and changed the corresponding BFI contrast only modestly; the response remained positive and the group-level flow result was unaffected.

**Figure S2.**
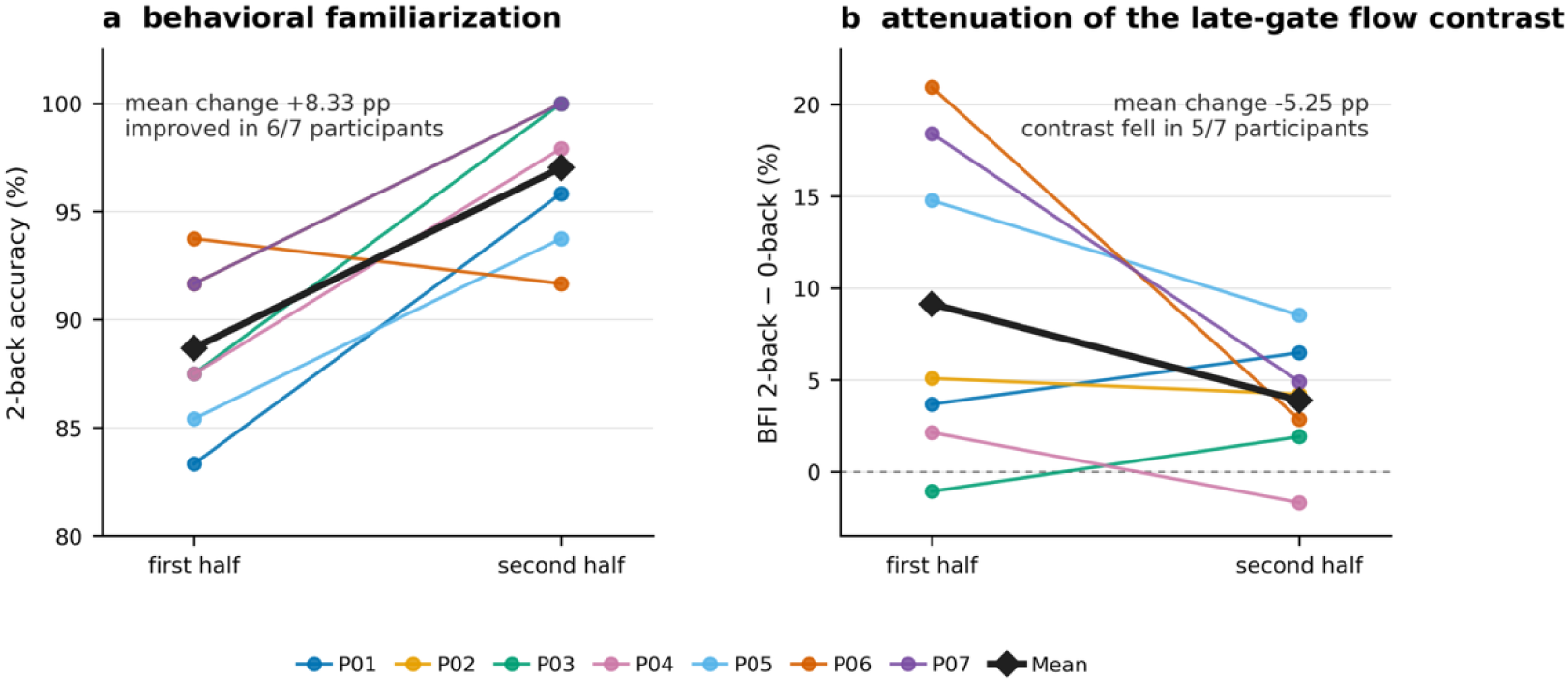
Behavioral familiarization and attenuation of the late-gate flow contrast (N=7). **a.** Mean 2-back block accuracy in the first and second halves of the session. Accuracy increased in six of seven participants, from a cohort mean of 88.7% to 97.0% (mean paired change, +8.33 percentage points). **b.** Primary 500–600-ps BFI 2-back minus 0-back contrast across the same session halves. The contrast decreased in five of seven participants, from a cohort mean of 9.13% to 3.89% (mean paired change, −5.25 percentage points; median paired change, −3.82 percentage points). Colored lines connect individual participants; black diamonds indicate cohort means. These trajectories are descriptive and do not establish behavioral–flow coupling or causality.

### S2. Stimulus–acquisition synchronization

The mobile and desktop applications communicated over a local network using connectionless UDP datagrams (port 55555). Messages were UTF-8 JSON carrying the event type, a unique session identifier, the mobile-device UTC timestamp, an optional time relative to session onset, and event-specific metadata (trial index, block, stimulus type, duration and participant response). The desktop application accepted events only after a session_start marker and rejected packets from a different session identifier. session_start triggered acquisition and session_end terminated it; for temporal alignment the software retained both the mobile-device timestamp and the desktop-receipt UTC time. All markers and their timing relative to acquisition onset were stored with the measurement data in a JSON manifest and embedded in the processed-session metadata.

**Table S1.** Summary of participant-level endpoints. Confidence intervals are participant-level intervals for the reported group effects. Forearm decomposition rows report paired Early–Late gate differences in the direction indicated. Values summarize analyses reported in the main text and introduce no additional statistical tests. P values are unadjusted.

| Experiment | Endpoint | Window / comparison | Estimate | 95% CI of effect | Inference |
| --- | --- | --- | --- | --- | --- |
| Forearm | Readout x TOF interaction | Same cuff window, Late / Early, wavelength-averaged | Median 2.61 octaves; 11/11 positive | 2.44 to 3.22 octaves | Exact two-sided P = 0.00098; primary |
| Forearm | BFI, 780 nm | Mean cuff suppression, Early to Late | 0.631 to 0.698 | Late-Early: 0.0404 to 0.0919 | Two-sided paired P = 0.00098; decomposition |
| Forearm | BFI, 852 nm | Mean cuff suppression, Early to Late | 0.583 to 0.618 | Late-Early: -0.0338 to 0.0953 | Two-sided paired P = 0.206; decomposition |
| Forearm | Delta $\mu_a$, 780 nm | Mean same-window response, Early to Late | 0.236 to 0.0397 cm<sup>-1</sup> | Early-Late: 0.123 to 0.280 cm<sup>-1</sup> | Two-sided paired P = 0.00195; decomposition |
| Forearm | Delta $\mu_a$, 852 nm | Mean same-window response, Early to Late | 0.218 to 0.0359 cm<sup>-1</sup> | Early-Late: 0.0996 to 0.274 cm<sup>-1</sup> | Two-sided paired P = 0.00195; decomposition |
| PFC | BFI, 780 nm | 500-600 ps, uncorrected | +5.07%; 7/7 positive | 1.46 to 8.68% | P = 0.014; exploratory primary readout |
| PFC | BFI + Early nuisance | 500-600 ps, joint GLM | +4.04%; 7/7 positive | 0.75 to 7.33% | P = 0.024; exploratory correction |
| PFC | BFI, rest-residualized | 500-600 ps | +3.69%; 6/7 positive | 0.25 to 7.13% | P = 0.039; sensitivity |
| PFC | Delta HbR | 525-625 ps | -2.84 x 10<sup>-3</sup> mM; 5/7 expected | -7.87 to 2.19 x 10<sup>-3</sup> mM | P = 0.22; secondary |
| PFC | Delta HbO | 525-625 ps | +3.19 x 10<sup>-3</sup> mM; 4/7 expected | -3.30 to 9.68 x 10<sup>-3</sup> mM | P = 0.27; secondary |

**Table S2.** Gate- and wavelength-resolved forearm photon support and BFI fit yield. Photon support is the gate-integrated baseline TPSF in native acquisition units, summarized across 12 recordings. BFI fit yield is the fraction of frames satisfying R ≥ 0.80. Rejected and total frame counts are pooled across recordings.

| Wavelength | Gate | Photon support, median [IQR] | BFI fit yield, median [IQR] | Rejected / total frames |
| --- | --- | --- | --- | --- |
| 780 nm | Early | 49,684 [41,760–56,946] | 100.0% [100.0–100.0%] | 2 / 1843 |
| 780 nm | Middle | 49,660 [41,384–57,178] | 100.0% [100.0–100.0%] | 1 / 1843 |
| 780 nm | Late | 19,359 [16,092–21,192] | 100.0% [100.0–100.0%] | 6 / 1843 |
| 852 nm | Early | 24,956 [20,852–28,253] | 100.0% [99.4–100.0%] | 11 / 1840 |
| 852 nm | Middle | 23,238 [19,997–26,551] | 100.0% [100.0–100.0%] | 7 / 1840 |
| 852 nm | Late | 8,233 [7,405–8,743] | 99.4% [96.1–100.0%] | 37 / 1840 |

**Table S3.** Robustness of the wavelength-averaged forearm readout-by-TOF interaction. The primary analysis used the final minute of stable occlusion before record-specific cuff release with standard BFI quality control and smoothing. Sensitivity analyses used the fixed 3.5–4.5-min interval, pre-censoring/pre-smoothing trajectories, or both. A two-wavelength interaction was considered evaluable only when both wavelength-specific response ratios were positive. One participant had an unstable uncensored 852-nm Late-gate BFI response in the raw sensitivity branch.

| Analysis | Evaluable / total | Median interaction (octaves) | Positive | Exact two-sided sign P |
| --- | --- | --- | --- | --- |
| Record-specific standard (primary) | 11 / 11 | 2.61 | 11 / 11 | 0.00098 |
| Record-specific, pre-censor/pre-smoothing | 10 / 11 | 2.55 | 10 / 10 | 0.00195 |
| Nominal 3.5–4.5 min, standard | 11 / 11 | 2.68 | 11 / 11 | 0.00098 |
| Nominal 3.5–4.5 min, pre-censor/pre-smoothing | 10 / 11 | 2.65 | 10 / 10 | 0.00195 |

